# Defining An Expanded RAS Conformational Landscape Based on Over 700 Experimentally Determined Structures of KRAS, NRAS, and HRAS

**DOI:** 10.1101/2022.02.02.478568

**Authors:** Mitchell I. Parker, Joshua E. Meyer, Erica A. Golemis, Roland L. Dunbrack

## Abstract

For many human cancers and tumor-associated diseases, mutations in the RAS isoforms (KRAS, NRAS, and HRAS) are the most common oncogenic alterations, making these proteins high-priority therapeutic targets. Effectively targeting the RAS isoforms requires an exact understanding of their active, inactive, and druggable conformations. However, there is no structure-guided catalogue of RAS conformations to guide therapeutic targeting or examining the structural impact of RAS mutations. We present an expanded classification of RAS conformations based on analyzing their catalytic switch 1 (SW1) and switch 2 (SW2) loops. From all 721 available human KRAS, NRAS, and HRAS structures in the Protein Data Bank (PDB) (206 RAS-protein complexes, 190 inhibitor-bound, and 325 unbound, including 204 WT and 517 mutated structures), we created a broad conformational classification based on the spatial positions of residue Y32 in SW1 and residue Y71 in SW2. Subsequently, we defined additional conformational subsets (some previously undescribed) by clustering all well modeled SW1 and SW2 loops using a density-based machine learning algorithm with a backbone dihedral-based distance metric. In all, we identified three SW1 conformations and nine SW2 conformations, each which are associated with different nucleotide states (GTP-bound, nucleotide-free, and GDP-bound) and specific bound proteins or inhibitor sites. The GTP-bound SW1 conformation can be further subdivided based on the hydrogen (H)-bond type made between residue Y32 and the GTP γ-phosphate: water-mediated, direct, or no H-bond. Further analyzing these structures clarified the catalytic impact of the G12D and G12V RAS mutations, and the inhibitor chemistries that bind to each druggable RAS conformation. To facilitate future RAS structural analyses, we have created a web database, called Rascore, presenting an updated and searchable dataset of human KRAS, NRAS, and HRAS structures in the PDB, and which includes a page for analyzing user uploaded RAS structures by our algorithm (http://dunbrack.fccc.edu/rascore/).

**Significance:** Analyzing >700 experimentally determined RAS structures helped define an expanded landscape of active, inactive and druggable RAS conformations, the structural impact of common RAS mutations, and previously uncharacterized RAS-inhibitor binding modes.

## INTRODUCTION

Mutations in the RAS isoforms, KRAS (4A and 4B splice forms), NRAS, and HRAS, drive oncogenesis in ~20% of human cancers, and cause a variety of tumor predisposition syndromes, making these proteins high-priority therapeutic targets (1). Over the past 30 years, our molecular understanding of RAS mutations and our ability to drug these proteins has considerably improved, owing, in part, to hundreds of structural studies examining wild type (WT) and mutated RAS in complex with various signaling effector and regulatory proteins, or with small-molecule and designed protein inhibitors (2). However, our structural understanding of RAS mutations is incomplete, and, except for KRAS G12C, G12D, and G13C, all mutated RAS forms have not yet been selectively targeted by therapeutics (3).

The RAS proteins are molecular switches that modulate growth and other signaling pathways in almost all cells of the human body by conformationally cycling between GDP-bound (“OFF”) and GTP-bound (“ON”) states (4). In normal tissues, RAS conformational cycling is tightly regulated by the catalytic CDC25 domain of guanine exchange factors (GEFs; e.g., SOS1), which remove GDP allowing subsequent GTP rebinding (5), and GTPase activating proteins (GAPs; e.g., NF1), which catalyze the otherwise slow intrinsic rate of GTP hydrolysis to GDP (6). GEFs and GAPs interact with the conformationally dynamic RAS switch 1 (SW1) and switch 2 (SW2) loops on RAS structures, which additionally provide binding interfaces for signal effector proteins (e.g., RAF1) and direct RAS inhibitors (7). RAS targeted therapies mostly bind to an SW1/SW2 pocket (called here “SP12”) to block RAS-protein interactions or an SW2 pocket (called here “SP2”) to lock RAS in an inactive, GDP-bound conformation. Overall, the configurations of SW1 and SW2 are essential to RAS function and the druggability of these proteins.

Most tumor-associated RAS mutations modify SW1 and SW2 conformational preferences and/or dynamics in ways that reduce the rate of intrinsic and GAP-mediated hydrolysis (residues 12, 13, and 61) and/or enhance the rate of GEF-mediated exchange (residues 13, 61, and 146) (8–12). The net effect of these alterations is to increase the steady-state cellular concentration of active, GTP-bound RAS that is capable of stimulating signaling pathways via the following mechanisms: by binding to and activating RAS effectors; by binding to the allosteric REM domain of the GEF SOS1, which functions to accelerate GDP release at the CDC25 domain of SOS1 (13); and by promoting homodimerization of RAS monomers at their helices α4 and α5, which is required for activation of certain dimeric effectors, such as RAF1 (14,15). However, the exact SW1 and SW2 conformations that can form each RAS complex and their potential druggability are unknown, making it difficult to design therapeutics that selectively block the activities of all WT and mutated RAS forms. Furthermore, for the most part, the described structural impact of many mutated RAS forms is underdetermined, either based on comparison of one or two mutated structures or extrapolations made while observing WT structures, necessitating further structural examination of RAS mutations.

While all published RAS structures have been made publicly available through the Protein Data Bank (PDB), this structural dataset has not been leveraged in a comprehensive way to improve our understanding of RAS conformations and mutations and to inform RAS drug discovery. Therefore, we analyzed all 721 available human KRAS, NRAS, and HRAS structures in the PDB to define a more comprehensive classification of active, inactive, and druggable RAS conformations and identify the structural consequence of common mutations. We first annotated the molecular contents of each RAS structure, including their mutation status, nucleotide state and bound protein (e.g., effector, GAP, GEF) or inhibitor site (e.g., SP12, SP2). Second, we conformationally classified all RAS structures based on the spatial positions of residue Y32 in SW1 and residue Y71 in SW2 and by the conformations of their catalytic SW1 and SW2 loops, as expressed by their backbone dihedral angles. By associating the identified SW1 and SW2 conformations with the annotated molecular contents of each structure, we were able to create a biologically and therapeutically informed map of the RAS conformational landscape and determine the structural impact of some common RAS mutations. Overall, our study expands our knowledge of RAS biology and provides a valuable resource for analyzing RAS structures in ways that will improve our understanding of RAS mutations and our ability to drug these proteins. The results of this work are presented in a continually updated database called Rascore (http://dunbrack.fccc.edu/rascore/).

## MATERIALS AND METHODS

### Preparing RAS structures

PDB entries containing human KRAS (all are the 4B isoform), NRAS, and HRAS were identified by SwissProt identifier in the *pdbaa* file (March 1 2022) on the PISCES webserver (http://dunbrack.fccc.edu/pisces/) (16). For each PDB entry, the asymmetric unit and all biological assemblies were downloaded and renumbered according to UniProt scheme using PDBrenum (http://dunbrack.fccc.edu/PDBrenum/) (17). In addition, electron density of individual atom (EDIA) scores (a per atom measure of model quality) (18) were downloaded from the ProteinPlus webserver (https://proteins.plus) (19).

Since some PDB entries contain multiple RAS polypeptide chains, each RAS chain of the asymmetric unit (only first model for NMR structures) was separated with its corresponding bound ligands and/or proteins. Ligands were labeled *Nucleotide*, *Ion*, *Inhibitor*, *Chemical*, *Modification*, or *Membrane* using a custom dictionary prepared in considering annotations from FireDB (https://firedb.bioinfo.cnio.es) (20). Since ligands were not always labeled by the RAS chain that they contact, all ligands were reassigned based on the following criteria:

1. *Nucleotide, Ion, Chemical, of Modification* – Possess the same chain label.
2. *Inhibitor* – Have more than 5 residue contacts within 4 Å of the chain.
3. *Membrane* – Link the chain to a nanodisc (synthetic membrane).

Proteins were assigned to a RAS chain based on the rules below with assignments checked against available biological assemblies and discrepancies corrected:

1. If they had more than 5 Cβ contacts within 12 Å and 1 atom contact within 5 Å of the RAS chain.
2. If they had more than 5 atom contacts within 5 Å of the RAS chain.
3. If they were the bounding protein component of a nanodisc.

Each RAS chain was treated as a unique RAS structure in subsequent analyses. Finally, RAS structures were annotated by various molecular contents, many of which are not reported in PDB entries (details provided in Supplementary Materials and Methods): mutation status, nucleotide state, bound protein, inhibitor site, inhibitor chemistry, and homodimer status. In addition, KRAS-NF1, KRAS-RASA1, HRAS-NF1, and HRAS-RASA1 co-complexes were modeled with the AlphaFold-Multimer software (21,22) within the ColabFold framework (23) utilizing the RASK_HUMAN (P01116-2), RASH_HUMAN (P01116), NF1_HUMAN (P21359-1, residues 1235-1451, RasGAP domain), and RASA1_HUMAN (P20936-1, residues 748-942, RasGAP domain) sequences.

### Conformational clustering

RAS structures were first subdivided into broad structural groups based on the spatial positions of residue Y32 in SW1 and residue Y71 in SW2. For Y32, two broad groups were defined:

1. *Y32in* – Y32 “in” the active site; Y32(OH)-G12(CA) distance < 10.5 Å.
2. *Y32out* – Y32 “out” of the active site; Y32(OH)-G12(CA) distance > 10.5 Å.

For Y71, two broad groups were defined:

1. *Y71in* – Y71 buried “in” the hydrophobic core; Y71(OH)-V9(CA) distance < 8.75 Å.
2. *Y71out* – Y71 facing “out” to the solvent; Y71(OH)-V9(CA) distance > 8.75 Å.

To identify conformational subsets within these spatial classes, for each nucleotide state, structures labeled Y32in and Y32out were clustered separately by the backbone configuration of their SW1 loops (residues 25–40), and similarly structures labeled Y71in and Y71out were clustered separately by the backbone configuration of their SW2 loops (residues 56–76).

In each conformational clustering, completely modeled loop structures possessing carbonyl (O) atom EDIA scores > 0.4 (indicating atoms well placed within the electron density) were clustered using the Density-Based Spatial Clustering of Applications with Noise (DBSCAN) algorithm with a backbone dihedral-based distance metric (previously implemented in refs. (24–27)). DBSCAN finds major clusters and removes outliers (28), which is ideal for clustering structural datasets since they usually contain several outliers that were poorly modeled or solved under rare experimental conditions. In this study, a distance metric was used that locates the maximum angular difference (*d*) upon pairwise comparison of the backbone dihedral angle values phi (*φ*), psi (*ψ*), and omega (*ω)* for residues 1 through *n* of compared loops *i* versus *j*, where *d*(*θ*_*i*_, *θ*_*j*_) = *2(1 – cos*(*θ*_*j*_ – *θ*_*i*_)):

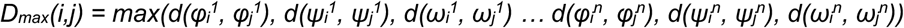

As in our previous study (24), DBSCAN was run across a grid of parameters and a set of quality control filters were applied to generate a robust consensus clustering (Supplementary Materials and Methods).

### Detecting hydrogen bonds

Hydrogen (H)-bond cutoffs were based on a previous analysis of protein structures in the PDB (29):

1. *H-bond* - 2.0-3.2 Å donor-acceptor distance with 90-180° carbon-donor-acceptor and carbon-acceptor-donor angles.
2. *Water-mediated (WM) H-bond*

a. 2.0-3.0 Å donor-water and acceptor-water distances with 80-140° carbon-water-acceptor and carbon-water-donor angles.
b. 2.8-5.6 Å donor-acceptor distance (arrived at using the Law of Cosines with the previously specified cutoffs and examining the distance distributions of WM H-bonds across RAS structures in the PDB).

### Data availability

All data are available as a static table in Supplementary Datasets S1-S4. In addition, our database called Rascore presents a continually updated dataset of all annotated and conformationally classified RAS structures in the PDB (http://dunbrack.fccc.edu/rascore/). The Rascore database includes a page for conformationally classifying user uploaded structures. Open-source code can be found in GitHub (https://github.com/mitch-parker/rascore).

## RESULTS

### Conformationally classifying RAS structures

We identified 721 human KRAS (N=436), HRAS (N=275), and NRAS (N=10) structures from 408 PDB entries (some entries contain multiple copies of the RAS protein, sometimes in different conformations). In all, there were 206 RAS-protein co-complexes, 190 inhibitor-bound, and 325 unbound structures, comprising 204 WT and 517 mutated structures (Supplementary Dataset S1). Subsequently, we created an automated system for annotating RAS structures by various molecular contents, including their mutation status, nucleotide state (“3P” for GTP or any triphosphate analog, “2P” for GDP, or “0P” for nucleotide-free), bound protein (effector, GAP, GEF CDC25 or REM domain, designed protein “binder,” or synthetic membrane “nanodisc”), small molecule inhibitor site (SP12 or SP2), and whether the α4α5 homodimer is present in the protein crystal (only X-ray structures). In this work, we defined an expanded RAS conformational classification by identifying the observed SW1 and SW2 conformations within the prepared dataset of RAS structures and associated each conformation with the annotated molecular contents to gain novel insights into WT and mutated RAS function and inhibition.

Several RAS conformations have been previously described based on the arrangement of SW1 and SW2. For SW1, the conformations are named by their nucleotide state and include: GDP-bound, nucleotide-free, and GTP-bound “state 1” (inactive) and “state 2” (active) (2); these conformations have been visually differentiated by the position of residue Y32 in SW1 relative to the active site (“in” or “out”) (**Fig. 1A**). For SW2, two GTP-bound conformations have been characterized, including an inactive, “T” state” (also called “off/ordered off” (30) and “state 2*” (31)) and an active, “R” state” (also called “on” (30)), which possess Y71 facing “out” to the solvent or buried “in” the hydrophobic core, respectively (**Fig. 1B**). Furthermore, other unnamed GDP-bound SW2 conformations have been differentiated based on their ability to bind certain small molecule inhibitors (32–34). However, only a few RAS structures in the PDB have been visually classified into the previously named and unnamed SW1 and SW2 conformations, and there is no systematic method for differentiating all known RAS conformations from each other and from potentially unidentified ones.

**Figure 1.**
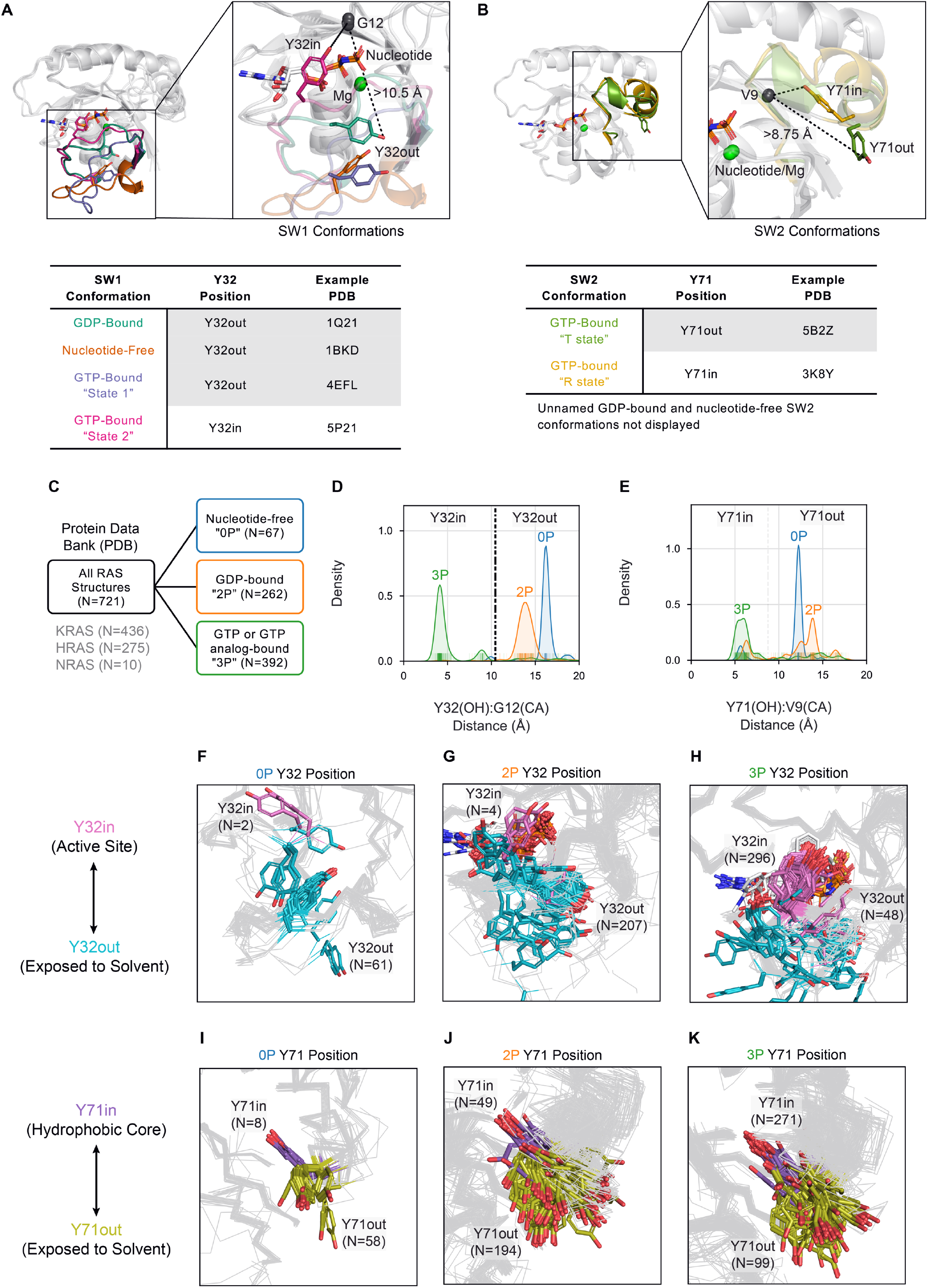
Broad structural classification of RAS structures. Previous conformational schemes based on the spatial position of, **A**, Y32 in SW1 and, **B**, Y71 in SW2. **C**, Separation of available RAS structures in the Protein Data Bank by nucleotide states 0P (nucleotide-free), 2P (GDP-bound), and 3P (GTP or GTP analog-bound). Distribution of distances by nucleotide state between, **D**, the hydroxyl (OH) atom of residue Y32 and alpha carbon (CA) atom of residue G12, and, **E**, the OH atom of residue Y71 and CA atom of residue V9; vertical dividing lines in plots indicate distance cutoffs for “in” versus “out” positions of Y32 (**D**) and Y71 (**E**), respectively. Structures classified Y32in (pink) and Y32out (cyan) within, **F**, 0P, **G**, 2P, and, **H**, 3P nucleotide states. Structures classified Y71in (purple) and Y71out (olive) within, **I**, 0P, **J**, 2P, and, **K**, 3P nucleotide states.

After separating RAS structures by 0P (N=67), 2P (N=262), and 3P (N=392) nucleotide states (**Fig. 1C**), we broadly classified structures into the known conformational scheme based on the spatial positions of residue Y32 relative to the active site (“Y32in” or “Y32out”) in SW1 and residue Y71 relative to the hydrophobic core (“Y71in” or “Y71out”) in SW2. By examining the distribution of distances between the Y32 hydroxyl (OH) atom and carbon alpha (CA) atom of residue G12 in the active site (**Fig. 1D**, Supplementary Dataset S2), we determined that almost all 0P (96.8%; N=61 of 63 classified) and 2P (98.1%; N=207 of 211 classified) structures are Y32out, as defined by a distance cutoff of 10.5 Å, while 3P structures contain a mix of Y32in (86.0%; N=296 of 344 classified; includes structures labeled state 2) and Y32out (14.0%; N=48 of 344 classified; includes structures labeled state 1) (Supplementary Table S1). The distribution of distances between the Y71 OH atom and CA atom of V9 in the hydrophobic core demonstrate that each nucleotide state contains a mix of Y71in and Y71out structures (defined by a distance cutoff of 8.75 Å), with 3P structures preferring Y71in and 0P and 2P structures preferring Y71out (**Fig. 1E**, Supplementary Dataset S2); the Y71in.3P and Y71out.3P structures include those classified as the R state and T state in the literature, respectively. In all, we were able to spatially classify 85.7% (N=618 of 721) of structures by the positions of Y32 and 94.2% (N=679 of 721) by the position of Y71 (Supplementary Table S1).

Since considerable conformational variability was observed for SW1 and SW2 within the defined Y32 and Y71 spatial groups by nucleotide state (**Figs. 1F–1K**), we sought to identify further conformational subsets than previously described. To do so, we analyzed RAS structures by their SW1 and SW2 configurations based on the backbone dihedral angle values of these loops: φ (phi), ψ (psi), and ω (omega). Since RAS structures in the PDB displayed the most dihedral variability on the Ramachandran map (35) (φ versus ψ plot) in residues 25-40 (SW1) and residues 56-76 (SW2), we selected these residue ranges to analyze (Supplementary Fig. S1). After removing loop structures with incomplete modeling or poor electron density, we arrived at 542 SW1 (75.2% of 721 structures) and 423 SW2 (58.7% of 721 structures) loop structures for conformational clustering. In our analysis, we used the Density-Based Spatial Clustering of Applications with Noise (DBSCAN) algorithm (28), which clusters points with sufficient numbers of near neighbors and classifies the remainder as outliers. We employed a distance metric that locates the maximum backbone dihedral difference upon pairwise comparison of loop residues (previously implemented in refs. (24–27)). We first separated RAS structures by nucleotide state (0P, 2P, and 3P) and spatial class (Y32in/out for SW1 and Y71in/out for SW2) and subsequently clustered the conformations of SW1 and SW2 within each group using DBSCAN. We then assigned a small number of poorly or incompletely modeled loops to the clusters obtained from DBSCAN through a nearest neighbors (NN) approach.

Overall, we were able to conformationally cluster 69.2% (N=375 out of 542) of SW1 and 56.7% (N=240 of 423) of SW2 loops that passed completeness and electron density checks. In addition, we assigned conformational labels by our NN approach to 85 SW1 and 51 SW2 loop structures that were not included in the original clustering due to removal by quality filters. Finally, we labeled any unclassified loops “outlier” if they were completely modeled or “disordered” if they were incompletely modeled. In all, we identified three SW1 and nine SW2 conformations, each of which were found across multiple RAS isoforms, PDB entries, and crystal forms (CFs; entries with the same space group and similar unit cell dimensions and angles) (Supplementary Table S2, including the mean dihedral distance and loop α carbon atom root-mean-square deviation for each conformation), indicating that these configurations are conserved within the residue interactions of the protein structure and are not solely the product of crystal contacts.

For clarity and brevity in our expanded RAS conformational classification, we named each SW1 and SW2 conformation by its spatial class, nucleotide state, and a conformational label (written in all-capital letters). Correlated counts of SW1 and SW2 conformations are provided in **Table 1** and reported throughout the text. The SW1 conformations are labeled Y32in.3P-ON (GTP-bound state 2), Y32out.2P-OFF (GDP-bound), and Y32out.0P-GEF (nucleotide-free) (**Fig. 2A**). There was no structurally uniform cluster within Y32out.3P structures that could be called the GTP-bound state 1. The only nucleotide-free SW2 conformation was Y32out.0P-GEF (**Fig. 2B**). The GTP-bound SW2 conformations included Y71in.3P-R (R state) and Y71out.3P-T (T state), and two previously unclassified druggable conformations associated with inhibitors at the SP12 site, which we named Y71in.3P-SP12-A and Y71.3P-SP12-B (**Fig. 2C**). The four GDP-bound SW2 conformations are all named for their predominant binding partners, which consist of SP2 and SP12 inhibitors and protein binders: Y71out.2P-SP2-A, Y71out.2P-SP2-B, Y71in.2P-SP12, and Y71out.2P-BINDER (**Fig. 2D**). Except for Y71out.2P-SP2-A, the other clusters labeled with SP12 or SP2 include structures both with and without bound inhibitors (Supplementary Dataset S1). In **Figs. 2E** and **2F**, the results for our SW1 and SW2 conformational clustering (with NN assignments added) are displayed as Ramachandran maps per residue of each cluster.

**Table 1.** Correlation of SW1 and SW2 conformational clusters.

| Conformation Label | Y32out.0P-GEF | Y32out.2P-OFF | Y32in.3P-ON | SW1 Outlier | SW1 Disordered | All |
| --- | --- | --- | --- | --- | --- | --- |
| Y71out.0P-GEF | 23 |  |  | 29 |  | 52 |
| Y71out.2P-SP2-A |  | 26 |  | 1 |  | 27 |
| Y71in.2P-SP12 |  | 20 |  |  |  | 20 |
| Y71out.2P-SP2-B |  | 19 |  |  |  | 19 |
| Y71out.2P-BINDER |  | 12 |  | 1 |  | 13 |
| Y71in.3P-R |  |  | 94 | 3 |  | 97 |
| Y71in.3P-SP12-A |  |  | 30 | 1 |  | 31 |
| Y71out.3P-T |  |  | 20 |  |  | 20 |
| Y71in.3P-SP12-B |  |  | 12 |  |  | 12 |
| SW2 Outlier |  | 59 | 53 | 85 | 53 | 250 |
| SW2 Disordered |  | 32 | 60 | 25 | 63 | 180 |
| All | 23 | 168 | 269 | 145 | 116 | 721 |

**Figure 2.**
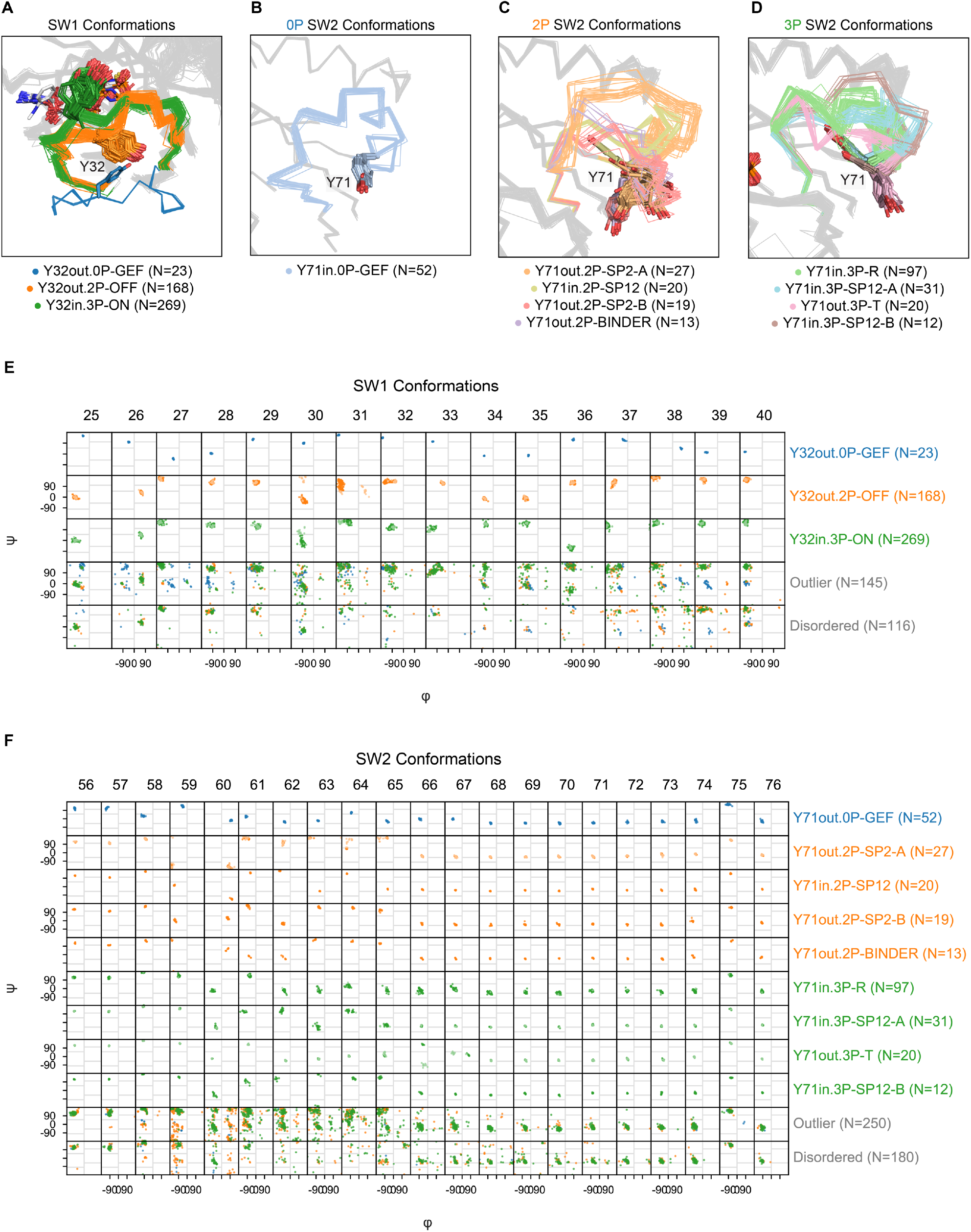
SW1 and SW2 conformational clusters. **A**, SW1 conformations. In the Y32out.0P-GEF conformation, the central Y32 residue in SW1 is ~12-13 Å from the active site. In the Y32out.2P-OFF conformation, SW1 is “closed” and interacts with the nucleotide through the backbone atoms of residues 28-32. In the Y32in.3P-ON conformation, further interactions are made with the nucleotide involving the side chains of residues Y32 and T35. SW2 conformations within, **B**, 0P, **C**, 2P, and, **D**, 3P states. In the Y71out.0P-GEF conformation, residues 58-60 of SW2 are pulled towards the nucleotide site, and the side chains of residues Q61 and Y71 form an intra-SW2 hydrogen bond (not displayed), which is not seen in other SW2 conformations. In all 2P SW2 conformations, except for Y71in.2P-SP12, Y71 is exposed to the solvent; the opposite trend is observed in all 3P SW2 conformations where Y71 is buried in the hydrophobic core of the protein, except in Y71out.3P-T where it is exposed. Ramachandran maps (φ versus ψ backbone dihedrals) for, **E**, SW1 and, **F**, SW2 conformational clusters. Lighter points on Ramachandran maps corresponding to loop structures with slight backbone differences from the remainder of the cluster.

### SW1 and SW2 conformations found in RAS-protein co-complexes

Currently, we do not know the combination of SW1 and SW2 conformations involved in each RAS interaction. Therefore, we analyzed which RAS conformations are found in the 206 RAS-protein complexes currently available in the PDB, except for nanodisc-linked structures which almost all were classified as outliers or disordered (**Table 2**).

**Table 2.** Distribution of SW1 and SW2 conformations by bound proteins.

| Conformation Label | Effector | GAP | GEF.CDC25 | GEF.REM | Binder | Nanodisc | Other | None | All |
| --- | --- | --- | --- | --- | --- | --- | --- | --- | --- |
| Y32out.0P-GEF |  |  | 23 |  |  |  |  |  | 23 |
| Y32out.2P-OFF |  |  |  |  | 13 |  | 1 | 154 | 168 |
| Y32in.3P-ON | 17 | 6 |  | 38 | 10 | 3 | 3 | 192 | 269 |
| SW1 Outlier | 5 | 1 | 32 |  | 11 | 10 | 3 | 83 | 145 |
| SW1 Disordered |  |  | 6 |  | 22 |  | 2 | 86 | 116 |
| Y71out.0P-GEF |  |  | 52 |  |  |  |  |  | 52 |
| Y71out.2P-SP2-A |  |  |  |  |  |  |  | 27 | 27 |
| Y71in.2P-SP12 |  |  |  |  |  |  |  | 20 | 20 |
| Y71out.2P-SP2-B |  |  |  |  |  |  | 1 | 18 | 19 |
| Y71out.2P-BINDER |  |  |  |  | 11 |  |  | 2 | 13 |
| Y71in.3P-R | 12 | 6 |  | 34 | 4 |  | 2 | 39 | 97 |
| Y71in.3P-SP12-A |  |  |  |  |  |  |  | 31 | 31 |
| Y71out.3P-T |  |  |  |  |  |  |  | 20 | 20 |
| Y71in.3P-SP12-B |  |  |  |  |  |  |  | 12 | 12 |
| SW2 Outlier | 8 | 1 | 9 | 4 | 40 | 11 | 4 | 173 | 250 |
| SW2 Disordered | 2 |  |  |  | 1 | 2 | 2 | 173 | 180 |
| All | 22 | 7 | 61 | 38 | 56 | 13 | 9 | 515 | 721 |

As expected, the SW1 conformation, Y32out.0P-GEF, and SW2 conformation, Y71out.0P-GEF, only exist in structures bound to the GEF.CDC25 domain of SOS1, which has SW1 held open by a SOS1 region called the “helical-hairpin” (5) (**Figs. 3A** and **3B**). As a new finding, we identified that the SW1 conformation, Y32in.3P-ON, and SW2 conformation, Y71in.3P-R, bind to the GEF.REM domain of SOS1 (**Figs. 3C** and **3D**), effectors (**Figs. 3E** and **3F**, including RAF1, PI3K, RALGDS, and PLCε, among others), and the GAP NF1 (**Figs. 3 G** and **H**). Originally, Y32in.3P-ON and Y71in.3P-R were not suggested to bind to GAPs (rather an unidentified GTP-bound “state 3”) (36,37). Furthermore, we found that slight positional variations of Y32 in Y32in.3P-ON structures influence whether the catalytic GAP “arginine (R)-finger” (6) of NF1 can enter the RAS active site: when Y32 is within 4.5 Å of the GTP γ-phosphate (**Fig. 3G**, left), the R-finger is excluded from the active site and, when it moves ~2 Å further from the γ-phosphate (**Fig. 3G**, right), the R-finger can enter the active site to potentially interact with GTP; this latter observation was made in two previous studies (9,38). Modeling KRAS-NF1, KRAS-RASA1, HRAS-NF1, and HRAS-RASA1 co-complexes using the AlphaFold-Multimer software (21,22), we found that RAS in these co-complexes was predicted to be Y32in.3P-ON and Y71in.3P-R with Y32 slightly shifted outward enabling the R-finger to enter the active site and directly interact with GTP, as observed in a previously published HRAS-RASA1 transition state structure (PDB: 1WQ1) (6) (Supplementary Fig. S2). Similar variability in the Y32 position is present in effector-bound structures (**Fig. 3E**). Later, we examine the functional significance of this Y32 positional variability as it relates to intrinsic and GAP-mediated hydrolysis and the structural impact of RAS mutations.

**Figure 3.**
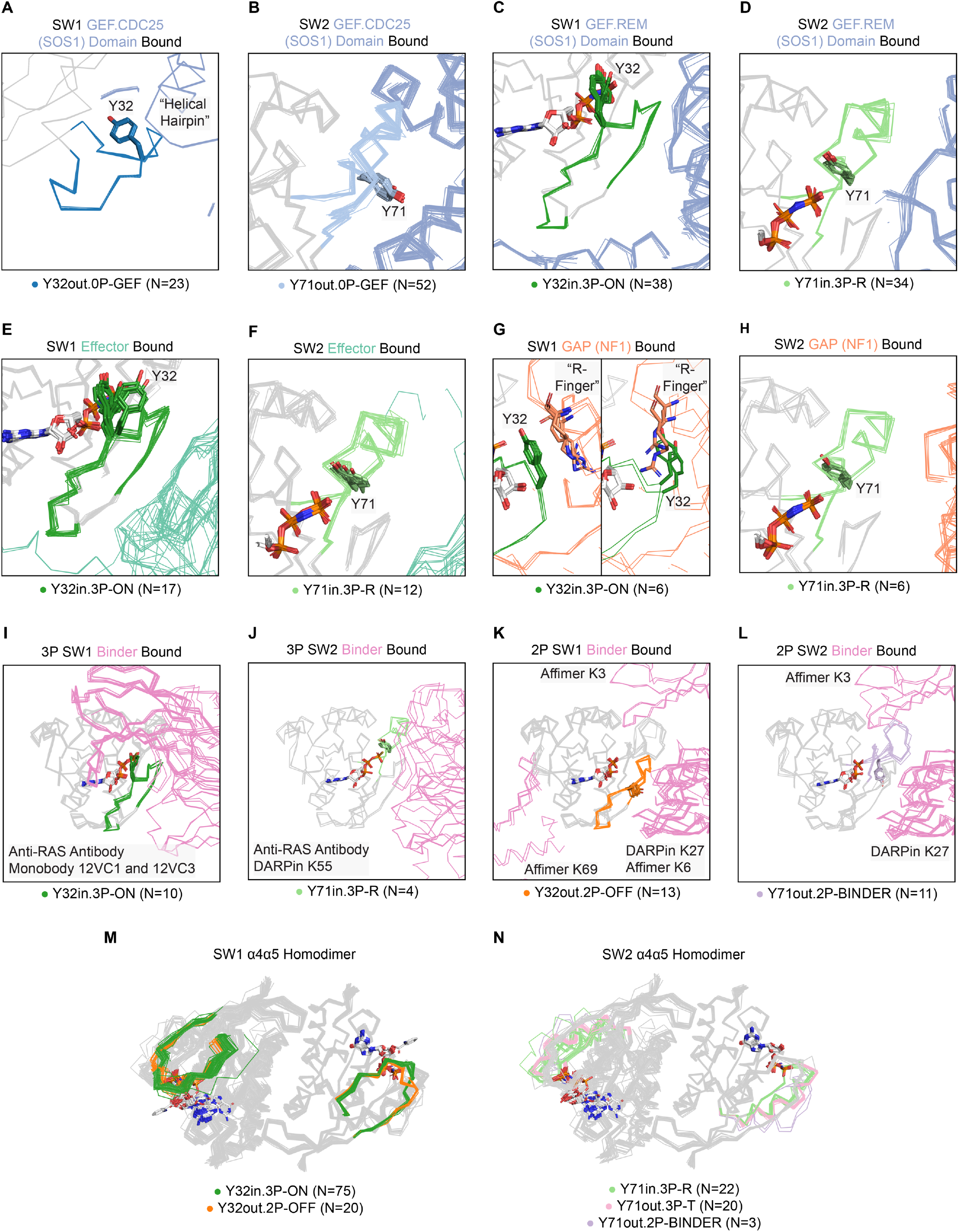
SW1 and SW2 conformations associated with bound proteins. SW1 and SW2 conformations bound to, **A** and **B**, the GEF.CDC25 (catalytic) domain of SOS, **C** and **D**, the GEF.REM (allosteric) domain of SOS1, **E** and **F**, effectors, **G** and **H**, the GAP NF1, **I** and **J**, 3P targeting designed protein “binders”, **K** and **L**, 2P targeting binders. **M** and **N**, structures forming the α4α5 homodimer. **A**, the “helical hairpin” of SOS1 opening SW1 of RAS. **G**, Comparison of catalytic “arginine (R) finger” position for the GAP NF1 with Y32 within 4.5 Å of the GTP γ-phosphate (left) and ~2 Å further away from the γ -phosphate (right, PDB: identical to the transition state stabilized in 1WQ1).

Besides the RAS co-complexes with signaling molecules described above, many RAS structures are found in the PDB bound to designed protein inhibitors (called here “binders”) (**Table 2**). In analyzing these co-complexes, we found that binders target three major sites on RAS structures: the SW1/SW2 pocket (SP12), the SW2 pocket (SP2), and the α4α5 interface (**Figs. 3I–3L**). In 3P structures, we identified that anti-RAS antibodies, the monobodies 12VC1 and 12VC3, and the DARPin K55 preferentially bind to the SW1 conformation, Y32in.3P-ON, and SW2 conformation, Y71out.3P-R, at the SP12 site. Since these loop conformations bind to signaling effectors as well (**Figs. 3I** and **3J**), it is no surprise that the listed binders are effective inhibitors of RAS-effector interactions (39–42). Next, we found that binders with preference for targeting 2P structures bind to multiple RAS interfaces; these include Affimers K3 (SP2 site), K6 (SP12 site), and K69 (α4α5 interface) as we all the DARPin K27 (SP12 site). Of the 2P-interacting binders, Affimer K3 and DARPin K27, which both function to block nucleotide exchange (41,43), bind to the SW1 conformation, Y32in.2P-OFF, and SW2 conformation, Y71out.2P-BINDER (hence the chosen conformational label) (**Figs. 3K** and **3L**). Moving forward, the identified binder interacting conformations can be used as structural templates in creating and optimizing additional protein inhibitors of RAS-effector interactions and nucleotide exchange.

### SW1 and SW2 conformations found in α4α5 homodimer complexes

Homodimerization of GTP-bound RAS monomers at their α4 and α5 helices is required in some cases for signal effector activation (e.g., RAF1) (14,15), and has been identified across numerous KRAS, NRAS, and HRAS crystal structures in the PDB (15,44). However, the conformations that can homodimerize are entirely unknown. We found the α4α5 homodimer across 144 HRAS (N=115), KRAS (N=28), and NRAS (N=1) structures (31% of X-ray experiments; 119 PDB entries and 19 crystal forms) (Supplementary Table S3). The functional relevance of the α4α5 homodimer is further supported by the observance of RAS α4α5 homodimers in co-crystal complexes with the signaling effectors RAF1 (N=5), PLCε1 (N=2), and RASSF1 (N=1) as well as the GEF, GRP4 (N=6) (Supplementary Dataset S1).

From the SW1 perspective, Y32in.3P-ON structures most commonly form the α4α5 homodimer (52.1%; N=75 of 144 dimers), with Y32out.2P-OFF forming this complex but less commonly (13.9%; N=20 of 144 dimers), and the remainder found in outlier or disordered structures (34.0%; N=49 of 144 dimers) (**Fig. 3M**). Surprisingly, we found that both Y71in.3P-R (active) and Y71out.3P-T (inactive) are the most common SW2 conformations (at approximately equal rates) that form the α4α5 homodimer (**Fig. 3N**), contrary to the expectation that only active RAS would form this complex.

### SW1 and SW2 conformations involved in inhibitor binding

RAS proteins are notoriously difficult to drug, because of their conformational variability and lack of deep surface pockets (3). Therefore, we sought to associate the presence of inhibitor-bound and -unbound sites with the set of identified SW1 and SW2 configurations, with the goal of cataloguing the potentially druggable RAS conformations of these proteins.

Utilizing the Fpocket software (45), we first obtained pocket descriptors for the observed inhibitor-bound sites on RAS structures, including their pocket volumes and druggability scores. Out of the 721 available RAS structures, 190 were inhibitor-bound: 44.7% at the SW1/SW2 pocket (SP12) site (N=85 of 190), 48.9% at the SW2 pocket (SP2) site (N=93 of 190), and the remaining 6.3% involving other or multiple sites (N=12 of 190) (**Table 3**) Subsequently, we focused our analysis on the most targeted pockets, SP12 and SP2. Overall, we were able to detect and calculate pocket descriptors for 92.9% of SP12 (N=79 of 85) and 91.4% of SP2 (N=85 of 93) inhibitor-bound sites (**Figs. 4A** and **4B**). We then used the Fpocket software to predict potentially druggable pockets in inhibitor-unbound structures and classified which were found at the SP12 or SP2 sites based on similarity of their residue contacts to observed inhibitor-bound sites. In sum, we identified 208 SP12 and 222 SP2 inhibitor-unbound sites, which translated to about 70% of these sites existing in the absence of inhibitor.

**Table 3.** Distribution of SW1 and SW2 conformations by inhibitor site.

| Conformation Label | SP12 |  | SP2 |  | Other |  | Multiple |  | All |
| --- | --- | --- | --- | --- | --- | --- | --- | --- | --- |
|  | Bound | Unbound | Bound | Unbound | Bound | Unbound | Bound | Unbound |  |
| Y32out.0P-GEF |  |  |  | 23 |  |  |  |  | 23 |
| Y32out.2P-OFF | 16 | 4 | 64 | 37 |  | 24 | 2 | 21 | 168 |
| Y32in.3P-ON | 53 | 97 | 1 | 24 |  | 65 | 3 | 26 | 269 |
| SW1 Outlier | 9 | 25 | 13 | 50 | 4 | 31 |  | 13 | 145 |
| SW1 Disordered | 7 | 12 | 15 | 18 | 3 | 51 |  | 10 | 116 |
| Y71out.0P-GEF | 1 |  |  | 48 | 3 |  |  |  | 52 |
| Y71out.2P-SP2-A |  |  | 26 |  |  |  | 1 |  | 27 |
| Y71in.2P-SP12 | 8 |  |  |  |  |  |  | 12 | 20 |
| Y71out.2P-SP2-B |  |  | 6 | 10 |  | 2 |  | 1 | 19 |
| Y71out.2P-BINDER |  | 1 |  | 8 |  | 4 |  |  | 13 |
| Y71in.3P-R | 11 | 37 |  | 7 |  | 34 |  | 8 | 97 |
| Y71in.3P-SP12-A | 14 | 11 |  | 2 |  |  |  | 4 | 31 |
| Y71out.3P-T |  | 11 |  | 1 |  | 6 |  | 2 | 20 |
| Y71in.3P-SP12-B | 7 | 3 |  |  |  | 2 |  |  | 12 |
| SW2 Outlier | 15 | 47 | 50 | 47 | 1 | 65 |  | 25 | 250 |
| SW2 Disordered | 29 | 28 | 11 | 29 | 3 | 58 | 4 | 18 | 180 |
| All | 85 | 138 | 93 | 152 | 7 | 171 | 5 | 70 | 721 |

**Figure 4.**
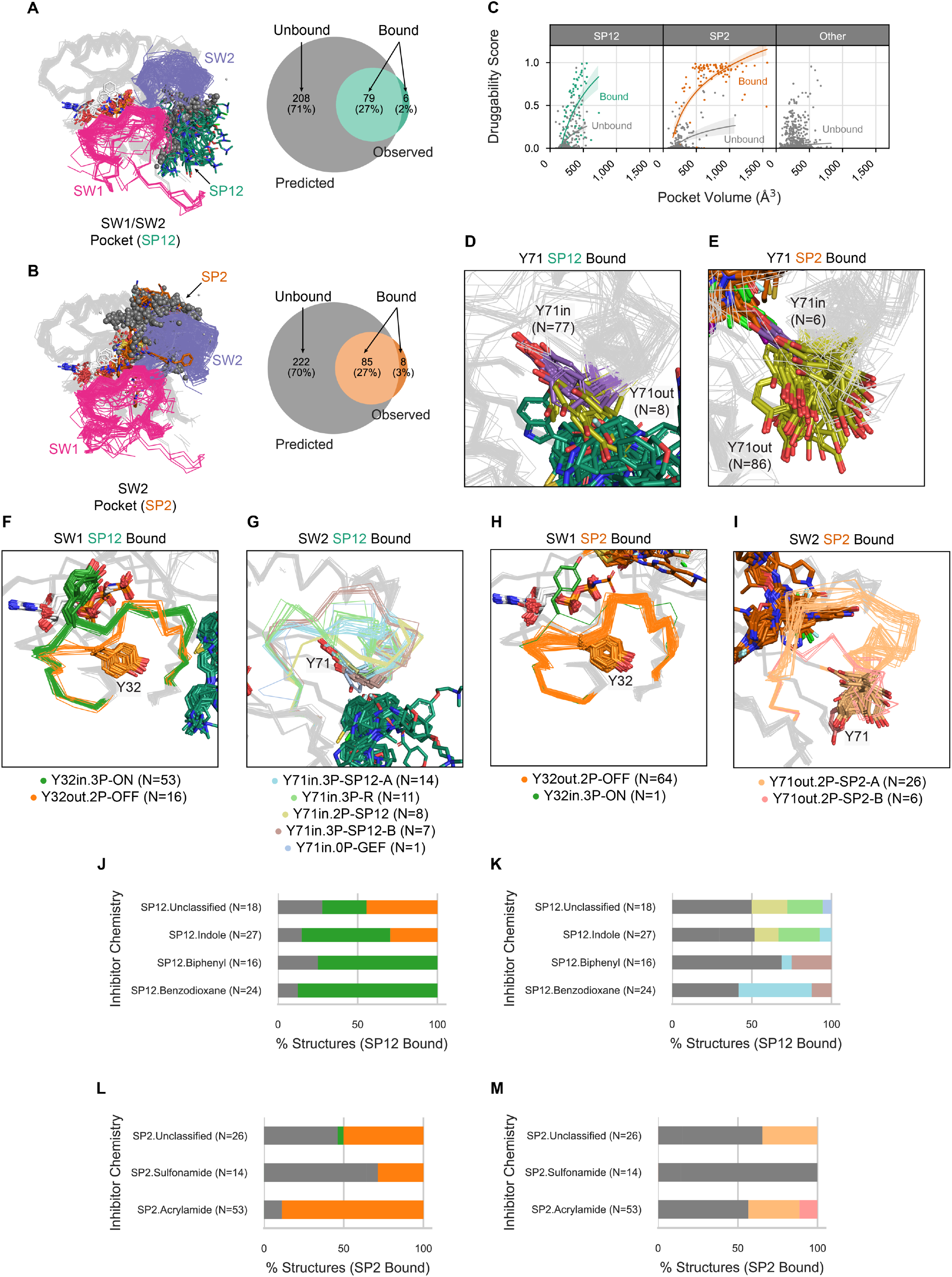
SW1 and SW2 conformations associated with inhibitor sites. Observed and predicted, **A**, SW1/SW2 pockets (SP12) and, **B**, SW2 pockets (SP2) across RAS structures in the PDB. **C**, Pocket volumes and druggability scores across inhibitor-bound and -unbound SP2, SP12, or other sites. Y71 positions in, **D**, SP12 and, **E**, SP2 inhibitor-bound structures. SW1 and SW2 conformations in RAS structures with, **E** and **F**, an inhibitor-bound SP12 site and, **G** and **H**, an inhibitor-bound SP2 site. Percent of each SW1 and SW2 conformation bound to inhibitors with different chemistries at, **I** and **J**, the SP12 site and, **K** and **L**, the SP2 site. **I-J**, colored by the same scheme as, **E-H**, with gray indicating structures labeled outlier or disordered.

Examining the calculated results for the SP12 and SP2 sites binding (Supplementary Dataset S3), we found that inhibitor-bound pockets correlated with higher druggability scores (>0.5) than inhibitor-unbound pockets (**Fig. 4C**). Other inhibitor-unbound sites were detected but with lower pocket volumes (<500 Å^3^) and druggability scores (<0.5) than the SP12 and SP2 sites. Importantly, we found that SP12 inhibitor-bound structures were mostly Y71in (90.2%; N=77 of 85), possessing an exposed SP12 site but an SP2 site occluded by the buried Y71 (**Fig. 4D**), while the SP2 inhibitor-bound structures by contrast were mostly Y71out (92.3%; N=86 of 93), with an exposed SP2 site but an SP12 site occluded by the exposed Y71 (**Fig. 4E**)

Further analyzing the conformational preferences of inhibitor-bound and -unbound structures, we found that SP12 inhibitors preferentially bind to the SW1 conformation, Y32in.3P-ON, and SW2 conformations, Y71in.3P-R, Y71in.3P-SP12-A, Y71in.3P-SP12-B, and Y71in.2P-SP12 (**Figs. 4F** and 4**G**). SP2 drugs alternatively bind to structures with the SW1 conformation, Y32out.2P-OFF, and SW2 conformations, Y32out.2P-SP2-A and Y32out.2P-SP2-B (**Figs. 4H** and **4I**). For both SP12 and SP2 sites, inhibitor-bound and -unbound structures had similar distributions of SW1 and SW2 conformations (**Table 3**). Notably, several SW1 or SW2 conformation that were found in both inhibitor-bound and -unbound structures correlated with the highest druggability scores (Supplementary Fig. S3), indicating that these inhibitor-binding conformations may be the most optimal targets for small molecule inhibition. In further support, no outlier conformations were found within the set of SP2 inhibitor-unbound sites with druggability scores >0.5, and only two structures were identified within the set of SP12 inhibitor-bound sites with scores greater than this cutoff (PDB: 1XCM and 4EFN, which are classified by some as GTP-bound state 1 (46)).

Recently, SP2 inhibitors with divergent core chemistries were found to bind to different SW2 configurations (32,33), but the conformational binding preferences of other SP2 and SP12 inhibitor chemistries are entirely unknown. We therefore subdivided SP2 and SP12 inhibitors by chemistries, focusing on inhibitor classes discussed repeatedly in the literature, and examined which SW1 and SW2 conformations each inhibitor chemistry binds to (Supplementary Table S4). The analyzed inhibitor chemistries included acrylamide and sulfonamide for SP2 inhibitors (34,47) and indole, benzodioxane, and biphenyl for SP12 inhibitors (48–53). SP12.Indole compounds preferred binding to structures with the SW1 conformation, Y32in.3P-ON and Y32out.2P-OFF, and SW2 conformations, Y71in.3P-R, Y71in.2P-SP12, or Y71in.3P-SP12-A (**Figs. 4J** and 4**K**); these SP12.Indole compounds included RAS-SOS1 inhibitors (e.g., DCAI) (54,55) and those that block multiple key RAS interactions (e.g., BI-2852, Cmpd2) (56). Most SP12.Indole compounds were found targeting KRAS G12D mutated structures in the PDB (Supplementary Dataset S1). In contrast, SP12.Benzodioxane and SP12.Biphenyl compound, which block key RAS-effector interactions (e.g., PPIN-1, PPIN-2) (49), were found to preferentially bind to structures with the SW1 conformation, Y32in.3P-ON, and SW2 conformations, Y71in.3P-SP12-A or Y71in.3P-SP12-B (**Figs. 4J** and **4K**); these inhibitors were mostly found targeting KRAS Q61H structures in the PDB (Supplementary Dataset S1). Lastly, SP2.Acrylamide compounds, which include the well-known KRAS G12C (covalent) inhibitors (e.g., sotorasib/AMG 510, adagrasib/MRTX849) (34,47), were found to preferentially bind to structures with the SW1 conformation, Y32out.2P-OFF, and SW2 conformations, Y71out.2P-SP2-A or Y71out.2P-SP2-B (**Figs. 4L** and **4M**). SP2.Sulfonamide compounds, which are another class of KRAS G12C (covalent) inhibitors, solely bound to structures labeled outlier or disordered.

### Structural impact of G12D and G12V mutations on intrinsic and GAP-mediated hydrolysis

Currently, there are at least 10 structures available in the PDB for each of the G12D, G12V, G12C, G13D, and Q61H mutated forms (Supplementary Dataset S1). Here we leverage our prepared dataset of RAS structures to elucidate a proposed theory regarding the active site configuration required for intrinsic and GAP-mediated hydrolysis and the structural consequence of the two most common RAS mutations, G12D and G12V on these activities.

Previously, Mattos and colleagues proposed that the G12D and G12V mutations may alter intrinsic hydrolysis by shifting the equilibrium between GTP-bound substates defined by the H-bond type made between the hydroxyl (OH) atom of Y32 and the closest γ-phosphate oxygen (called here O1G) atom of GTP or GTP analogs: one within direct hydrogen (H)-bonding distance, which they theorized to be catalytically incompetent, and another within water-mediated (WM) H-bonding distance, which they theorized to be catalytically competent (57,58). Given the previous observation of multiple Y32 positions in RAS-effector (**Fig. 3E**) and RAS-GAP (**Fig. 3G**) co-complexes, we wondered if there are possibly two or more hydrolytically relevant substates within the GTP-bound equivalent (i.e., 3P) structure. Examining the distribution of distances between the Y32(OH) and 3P(O1G) atoms (Supplementary Dataset S4), we found three peaks at distances of 3, 4.5, and 7 Å, which we associated with the observance of WM, direct, and no H-bonds, respectively (**Fig. 5A**).

**Fig. 5.**
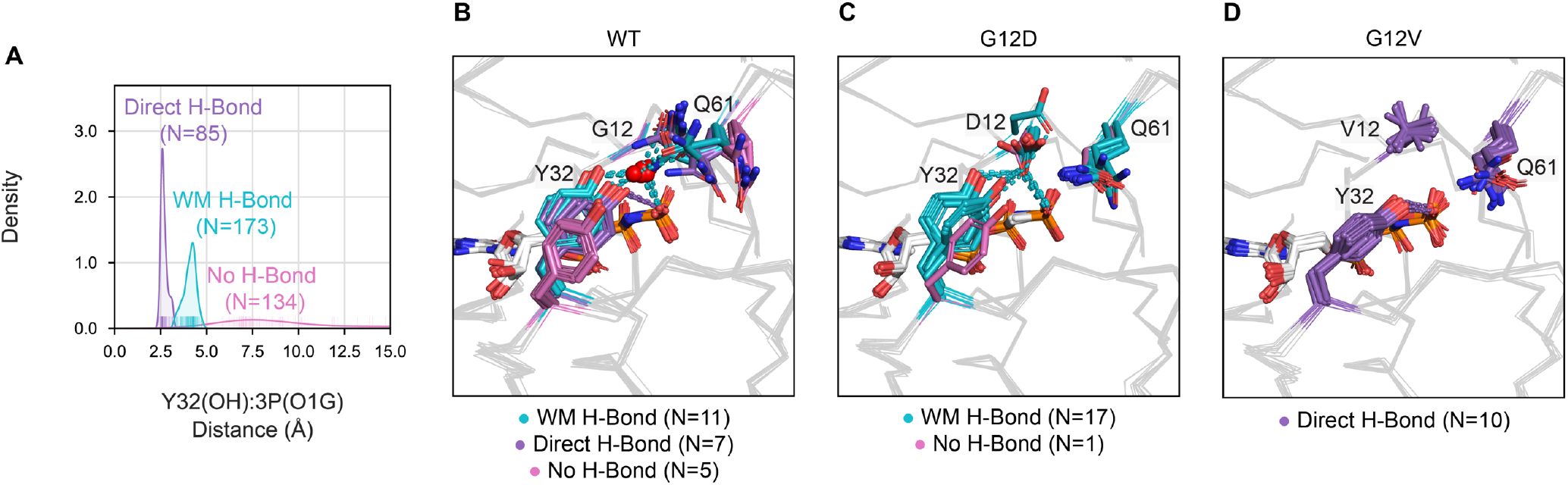
Structural impact of G12D and G12V mutations on GTP-bound substate preference. **A**, Distance distribution within 3P structure between the hydroxyl (OH) atom of residue Y32 and closest γ-phosphate (called here O1G) atom of GTP or GTP analogs, which was used to define hydrogen (H)-bonding subtypes: water-mediated (WM) H-bond, direct H-bond, and no H-bond. 3P substate preference within, **B**, WT, **C**, G12D, and, **D**, G12V structures.

Next, we compared the 3P substate preferences of WT, G12D, and G12V structures (included KRAS and HRAS isoforms) with the SW1 conformation, Y32in.3P-ON, and SW2 conformation, Y71in.3P-R, since these configurations define the active form of RAS proteins that occurs immediately before their GTP to GDP transition. While WT structures (N=23) had a nearly even distribution of 3P substates (**Fig. 5B**), 94% of G12D structures (N=17 of 18) were observed in the WM H-bond substate (**Fig. 5C**) and 100% of G12V structures (N=10 of 10) were found in the direct H-bond substate (**Fig. 5D**). The preference of G12V and G12D mutated structures for the WM H-bond (catalytically competent) and direct H-bond (catalytically incompetent) substates, respectively, may explain why G12V mutations severely impair intrinsic hydrolysis while G12D mutations only slightly dampen the hydrolytic reaction (8). Of note, we found that Y32 is stabilized in the WM H-bond substate in G12D mutated structures by residue D12 replacing the location of the water molecule (**Fig. 5C**), suggesting that intrinsic hydrolysis can still be preserved if Y32 is held within WM H-bonding distance of the GTP γ-phosphate. Relating the identified 3P substates to GAP-mediated hydrolysis, we propose that the preference of G12D and G12V mutated structures for substates other than no H-bond is the likely reason why GAP-mediated hydrolysis is almost non-existent in the context of these mutations (4).

## DISCUSSION

Since the first HRAS structure was experimentally solved in 1990, researchers have focused on characterizing the RAS conformational landscape by examining the possible structural configurations of their catalytic SW1 and SW2 loops. In this study, we used an extended dataset (721 KRAS, NRAS, and HRAS structures), and an approach that differs from previous studies (59,60), to create a data-driven RAS conformational classification, identifying three SW1 and nine SW2 RAS conformations. Our approach can be used to automatically conformationally classify and annotate the molecular contents of additional RAS structures as they are experimentally solved, providing a clear and consistent method for comparing WT and mutated structures across various biological and inhibitory contexts. To facilitate future RAS structural analyses, we have created a database, called Rascore, presenting the results from this study, which includes a page for conformationally classifying user uploaded RAS structures (http://dunbrack.fccc.edu/rascore/).

One uncertainty faced in defining our RAS conformational classification was identifying which GTP-bound SW1 conformations are state 1 and state 2. These SW1 conformations were discovered in the early 2000s with the observation of two peaks in the ^31^P NMR spectra for the GTP α and γ phosphates (61). Later studies found that mutations in residues Y32 and T35, as well as the common G12V mutation, cause a shift to the state 1-associated peaks, while other mutations, such as G12D, and the presence of the signaling effector, RAF1, cause a shift to state 2 (62–64). Subsequently, researchers experimentally solved a potential state 1 structure using a T35S mutant construct (65), and following for WT, G12V, Q61L, and other mutations (46,66). However, the previously labeled state 1 structures, which are found within our classified set of Y32out.3P structures, occurred too infrequently in this analysis to unambiguously call them the biological state 1 conformation. Moreover, the identified H-bonding substates within 3P structures (WM H-bond, direct H-bond, and no H-bond), could also explain the split state 1 and state 2 peaks in NMR spectra. Considering the NMR studies described above, and that we found G12D and G12V mutated structures prefer the WM H-bond and direct H-bond substates, respectively, we propose that the WM H-bond substate is state 2 and that state 1 is either the direct H-bond or no H-bond substates, some part of Y32out.3P, or some mixture of these structural configurations.

In contrast to other studies (59,60), we associated each SW1 and SW2 conformation with RAS interactions involving bound proteins and small molecular inhibitors. Overall, our analysis confirmed several previously held hypotheses regarding RAS conformations in a large experimental dataset and helped us uncovered some new hidden trends. For example, it has been hypothesized that RAS preferentially binds to signaling effectors and the GEF.REM domain of SOS1 when its SW1 conformation is “GTP-bound” (our Y32in.3P-ON) and its SW2 conformation is in the “R state” (our Y71in.3P-R) (13,30,31). We confirmed these observations and discovered that these same conformations bind to GAPs as well. In addition, we validated that both GTP-bound and GDP-bound SW1 structures can form the α4α5 homodimer, as was previously shown through NMR experiments of KRAS (14). Furthermore, we found that both the active, Y71in.3P-R and inactive, Y71out.3P-T SW2 conformations can α4α5 homodimerize, which was an observation not previously reported.

Another major value of this work is that it defined a comprehensive set of RAS conformations that are known targets for small molecule or designed protein inhibitors. Six out of seven druggable SW2 conformations are newly characterized in this study (all except for Y71in.3P-R); these include: GTP-bound Y71in.3P-SP12-A and Y71in.3P-SP12-B, and GDP-bound Y71in.2P-SP12, Y71out.2P-SP2-A, Y71out.2P-SP2-B, and Y71out.2P-BINDER. We associated each of these conformations with their preference for binding inhibitors with certain core chemistries, which is information researchers can use to select appropriate structural templates during structure-guided drug design. One overall finding from our analysis was that all the identified druggable RAS conformations exist in the absence of inhibitors, indicating that these structural configurations may naturally occur within a biological context and are not solely the product of an induced fit model. In addition, we found that the SP2 inhibitor site is mainly present in structures with Y71 exposed to the solvent (Y71out), while the SP12 inhibitor site appears in structures with Y71 buried into the protein core (Y71in). The consistency of this finding among many inhibitor-bound structures suggests it is an essential determinant of SP2 and SP12 druggability.

While this study has expanded our understanding of RAS structural biology, it only marks the beginning of characterizing the RAS conformational landscape. We hope that our RAS conformational classification system will be paired with further structure-activity relationship data to create machine learning models for RAS drug discovery. The pharmaceutical industry has over six times as many RAS inhibitor-bound structures as there are available in the PDB (67), and analysis of these structures using our conformational classification approach can help in identifying further druggable RAS conformations. Most importantly, having all RAS structures in the PDB consistently annotated and conformationally classified will enable simple utilization of this growing structural dataset for informing RAS drug discovery and studies of RAS mutations in human cancers and other diseases.

## Acknowledgments

We thank Bulat Faezov for providing the program PDBrenum in advance of publication, Vivek Modi for sharing scripts for calculating dihedral angles, and Simon Kelow for guidance in developing the conformational clustering algorithm.

## SUPPLEMENTARY MATERIALS AND METHODS

### Annotating RAS structures

RAS structures were annotated by various molecular contents:

1. *Mutation Status* – Identified by comparison of the sequence in each PDB entry to the human UniProt sequences for KRAS (P01116-1 and −2), NRAS (P01111), and HRAS (P01112).
2. *Nucleotide State –* The following labels were ascribed:

a. 0P – Nucleotide-free (i.e., none)
b. 2P – GDP-bound (only GDP)
c. 3P – GTP or GTP analog-bound (e.g., GTP, GppNHp, etc.)
3. *Bound Proteins –* Labeled by Pfam (68) or SwissProt identifier:

a. Effector – Pfams RBD, RA, and PI3K_rbd
b. GEF.CDC25 – Pfam RasGEF not bound to 3P nucleotide state
c. GEF.REM – Pfam RasGEF bound to 3P nucleotide state
d. GAP – Pfam RasGAP
e. Binder – No Pfam
f. Nanodisc – SwissProt APOA1_HUMAN
4. *Inhibitor Site* – Inhibitors were classified by binding site based on the presence of one or more predefined residue contacts within 4 Å of the RAS chain. Subsequently, inhibitor unbound sites were detected with Fpocket (45) and assigned to an inhibitor site if their average Simpson similarity (based on residue contacts) was greater than 0.6 to all inhibitor bound sites:

a. SP2 – 12, 96, or 99
b. SP12 – 5, 39, or 54
c. Other

i. Nucleotide Site – 29, 30, or 32
ii. Base of Nucleotide Site – 85, 118, or 119
iii. Allosteric Site (near C-terminal end) – 4, 49, or 164
iv. P110 Site (near residue P110) – 106, 108, or 110
5. *Inhibitor Chemistry –* The following SMILES strings were used in searching for inhibitor chemistries (performed with RDKit):

a. SP2

i. Acrylamide - CC(=O)N1CCNCC1 (1-Acetylpiperazine), CC(=O)N1CCC1 (1-Azetidin-1-YL-ethanone), or CC(=O)N1CCCC1 (N-Acetylpyrrolidine)
ii. Sulfonamide - CCS(=O)(=O)N (Ethanesulfonamide) or CC(=O)N1CCCCC1 (1-Acetylpiperidine)
b. SP12

i. Indole - C1=CC=C2C(=C1)C=CN2 (Indole)
ii. Benzodioxane - C1COC2=CC=CC=C2O1 (1,4-Benzodioxane)
iii. Biphenyl - C1=CC=C(C=C1)C2=CC=CC=C2 (Biphenyl)
6. *Homodimer Status –* The α4α5 homodimer was identified using the protocol employed in the Protein Common Interface Database (ProtCID) (44), requiring an average Q-score greater than 0.3 to the α4α5 homodimer found in PDB: 3K8Y.

### Conformationally Clustering SW1 and SW2

Well-modeled SW1 and SW2 loops were conformationally clustered using the DBSCAN algorithm with a backbone dihedral-based distance metric. Similar to a previous study (24), DBSCAN was run across a grid of parameters and a set of quality control filters was applied to generate a robust consensus clustering. This procedure was necessary since a single setting of DBSCAN cannot identify all possible conformational clusters due to their varying sizes, shapes, and densities. Below, are the detailed steps involved in our conformational clustering pipeline:

#### 1. Run DBSCAN on Well Modeled Loops

Different parameters of DBSCAN can produce slightly divergent clustering results with merging, splitting, or disappearance of clusters. Therefore, DBSCAN was run across a grid of parameters *D*=0.1-1.6 for ε (~20-80°) with steps of 0.1 and minimum samples 7-15 with steps of 1 and, following, a consensus of these clustering results was taken. The ε range covers the smallest regional subdivision of the Ramachandran map that residues with similar dihedrals can belong to (35). This range was found to be ideal for clustering in a previous study (24).

#### 2. Find Passing Clusters Across Runs

Not all DBSCAN clustering runs produce ideal separation of clusters. Therefore, two quality control filters we used to remove non-optimal clusters across runs before performing the consensus procedure: (a) mean silhouette score and (b) maximum dihedral distance:

a. Silhouette score is a measure incorporating the similarity of an object to members of its own cluster (cohesion) and difference from other clusters (separation) (69). The score can range from −1 (poor match to cluster) to 1 (good match to cluster). Entire clusters with mean silhouette score less than 0.6 were removed.
b. Some larger DBSCAN parameters can merge similar conformational clusters that are separate conformations. Therefore, clusters with a maximum dihedral distance greater than *D*=3.75 (~150°) were removed. Clusters with points this far apart tend to be a mix of two Ramachandran regions and, therefore, unsuitable for our purposes.

#### 3. Get Union of Similar Clusters Across Runs

Upon removal of poor clusters through quality filters, the union of clusters across runs with a Simpson similarity score greater than 0.9 were taken. The Simpson similarity score is the number of points two clusters have in common divided by the size of the smaller cluster. In most cases, across DBSCAN runs, a cluster found at smaller values of ε were often a subset of another cluster (i.e., Simpson score of 1.0) found at larger values of ε, as outlying points of the cluster were incorporated.

#### 4. Merge Clusters with Close Loop Cα-RMSD

The maximum dihedral metric is highly sensitive to peptide flips that an author may accidentally structurally model but does not signify a different conformation. This often happens at low resolution when the electron density can be modeled in two different ways.

Therefore, clusters with a loop Cα-RMSD less than 1.2 Å were merged, which was found to be an appropriate cutoff through trial and error, by testing a range of 0.5-2.0 Å and visualizing the results.

#### 5. Prune Cluster Members

At certain DBSCAN settings, some bordering structures can find their way into clusters that visually appear to be outliers. In consequence, cluster members were pruned if they had a nearest neighbor dihedral distance greater than *D*=0.45 (~40°), which covers half of the smallest regional subdivision of the Ramachandran map, or loop Cα-RMSD greater than 1.2 Å.

#### 6. Remove Small Clusters

Since a wide range of DBSCAN parameters are traversed during clustering, some small conformational clusters can be identified that have no functional or binding corollaries or are duplicates from a single study or set of experimental conditions. To filter out these unmeaningful conformations, clusters possessing less than seven chains or less than five PDB entries were removed.

#### 7. Classify Poorly Modeled Loops

Since the poorly modeled loops were not included in clustering a reversal of the pruning approach was used to assign these loops to clusters. Poorly modeled loops were only classified to clusters if their mean NN dihedral distance was less than 0.45 (~40°) or loop Cα-RMSD was less than 1.2 Å in reference to a single conformational cluster. This approach can be used to conformationally classify additional RAS structures that will be experimentally solved and deposited to the PDB, or ones produced through computational simulations. A similar dihedral distance cutoff was used in a previous study (70).

## SUPPLEMENTARY FIGURES AND TABLES

**Supplementary Figure S1.**
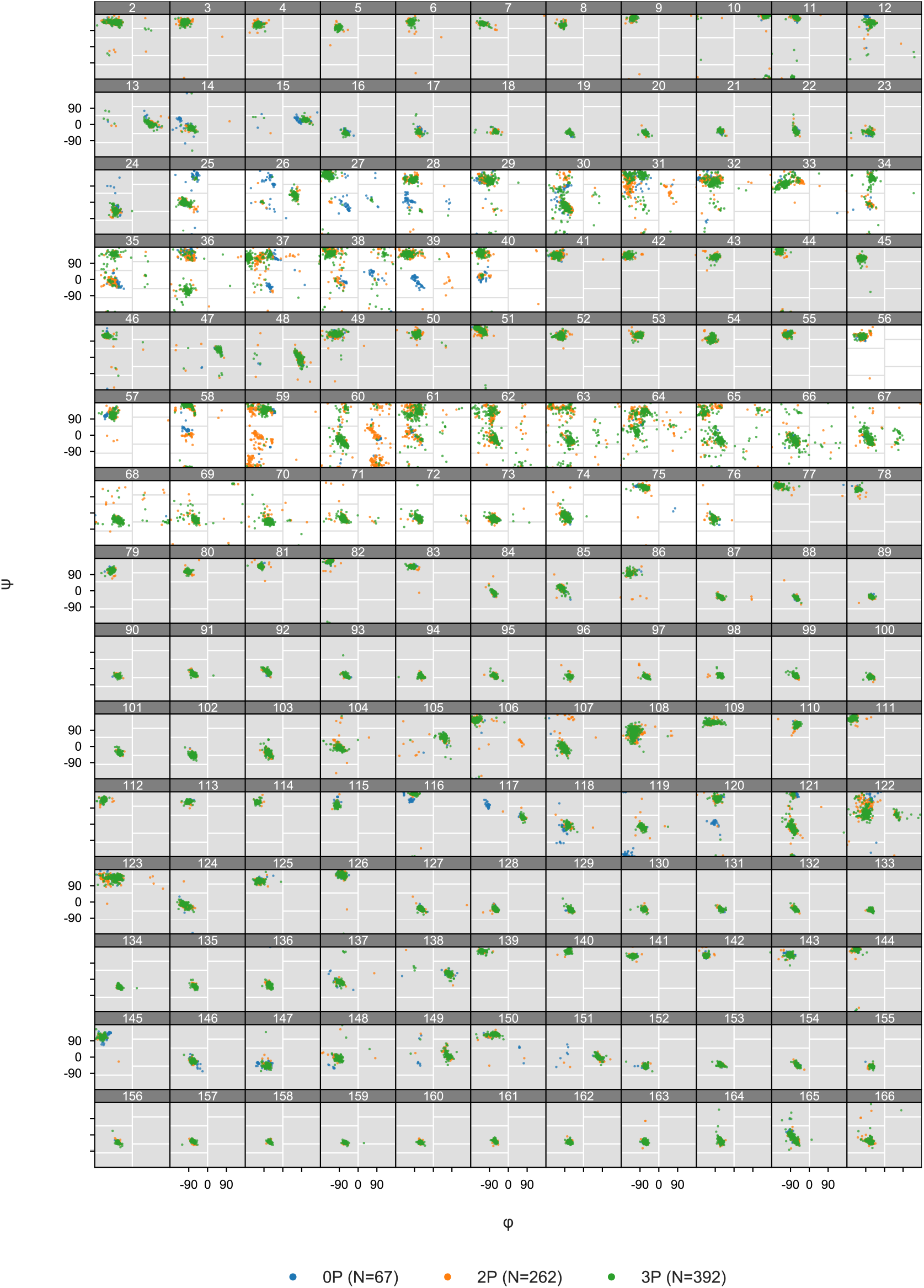
Ramachandran map (φ versus ψ backbone dihedrals) for available RAS structures in the PDB. Includes the RAS GTP-binding (G) domain (residues 1-166). SW1 (residues 25-40) and SW2 (residues 56-76) loops highlighted.

**Supplementary Figure S2.**
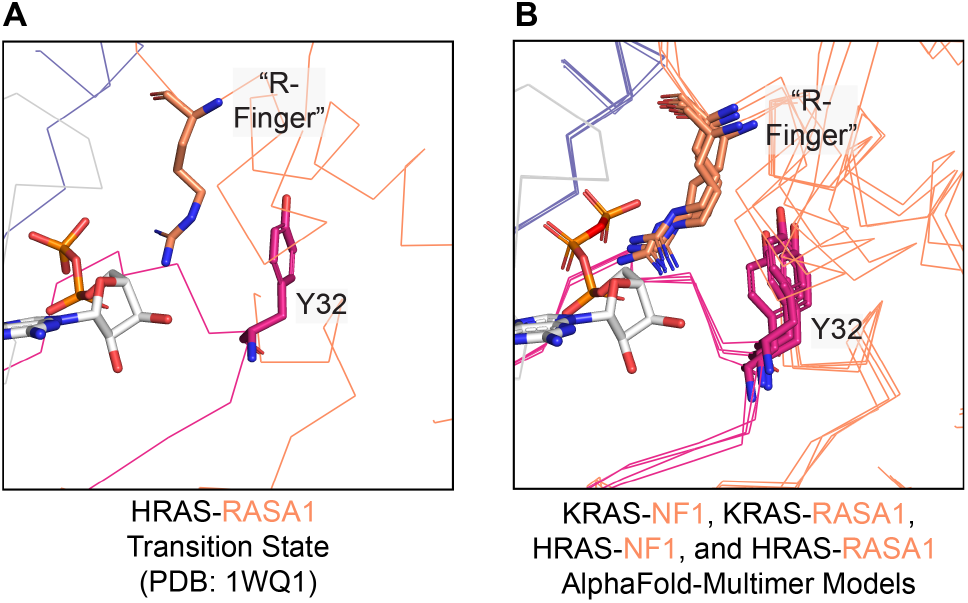
Comparison of experimentally determined and modeled RAS-GAP co-complexes. **A**, Transition state of GAP-mediated hydrolysis in an HRAS-RASA1 co-complex (PDB: 1WQ1). **B**, KRAS-NF1, KRAS-RASA1, HRAS-NF1, and HRAS-RASA1 co-complexes generated with AlphaFold-Multimer, which predicted the catalytic “arginine (R)-finger” to be within interacting distance to the GTP, similar to what is observed in the experimentally determined HRAS-RASA1 co-complex (**A**).

**Supplementary Figure S3.**
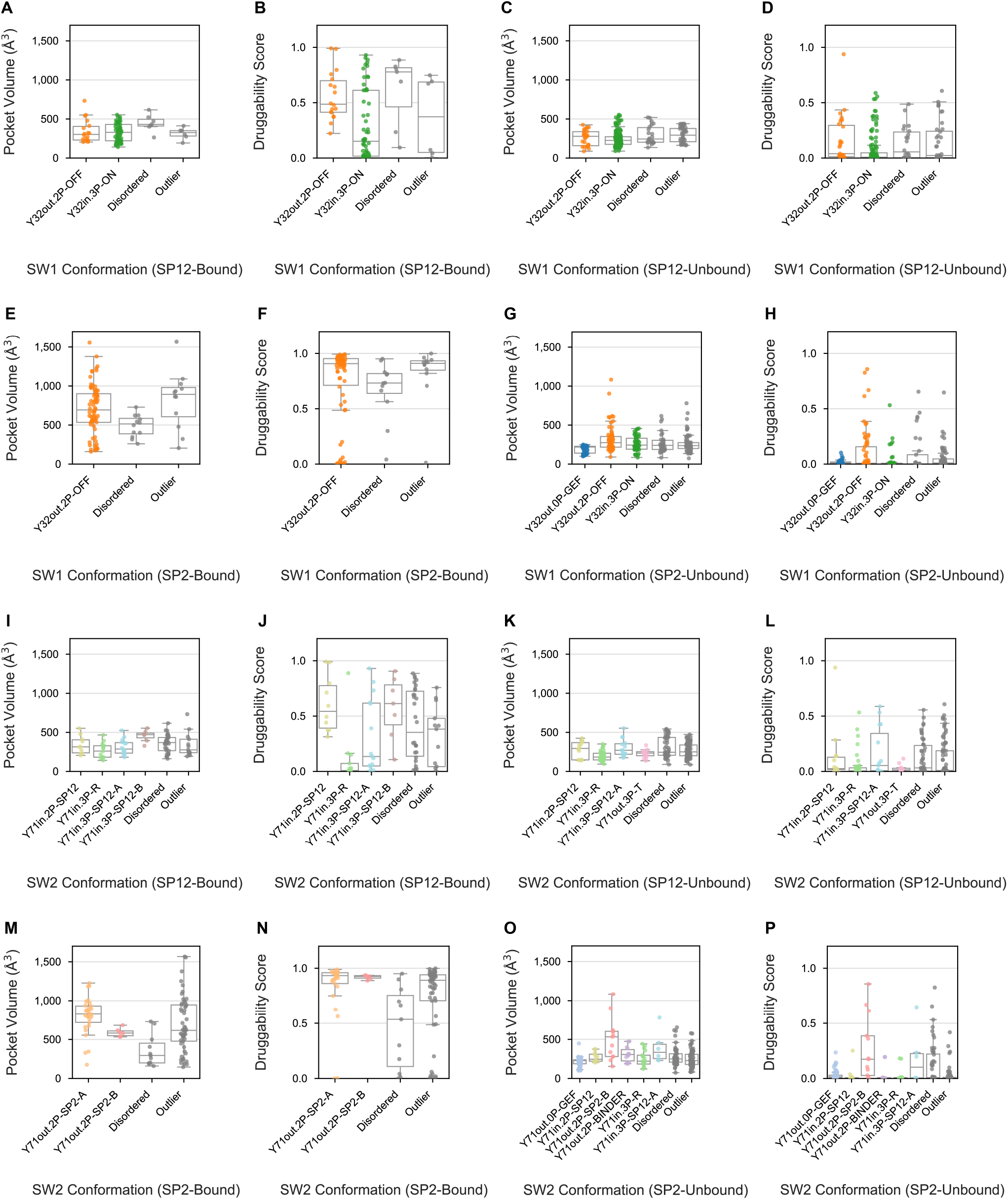
Pocket volumes and druggability scores for SP12 and SP2 inhibitor-bound and - unbound sites. Pocket volumes and druggability scores for SW1 conformations with SP12 inhibitor sites that are, **A** and **B**, bound and **C** and **D**, unbound; SW1 conformations with SP2 inhibitor sites that are, **E** and **F**, bound and **G** and **H**, unbound; SW2 conformations with SP12 inhibitor sites that are, **I** and **J**, bound and, **K** and **L**, unbound; and SW2 conformations with SP2 inhibitor sites that are, **M** and **N**, bound and, **O** and **P**, unbound. Only conformations with more than three structures with pockets are displayed.

**Supplementary Table S1.**
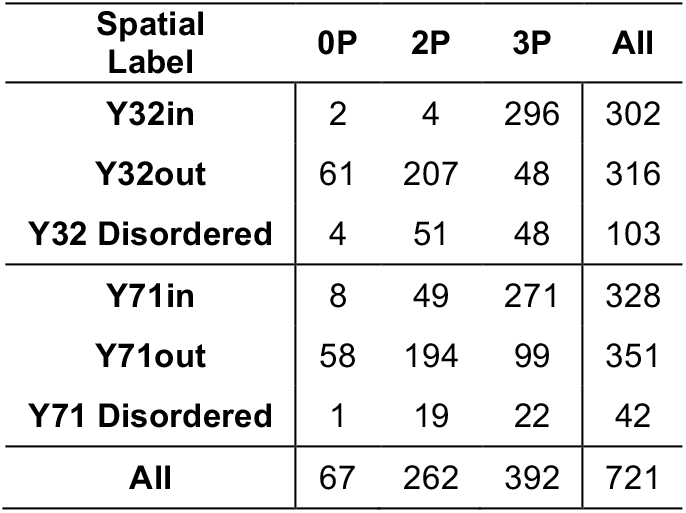
Distribution of Y32 and Y71 spatial positions by nucleotide state.

**Supplementary Table S2.** Summary of SW1 and SW2 conformational clusters.

| Conformation Label | Total Structures |  |  |  |  |  | NN Distance |  |
| --- | --- | --- | --- | --- | --- | --- | --- | --- |
|  | N | PDB | CF | HRAS | KRAS | NRAS | Backbone Dihedral (°) | Loop CA-RMSD (Å) |
| Y32out.0P-GEF | 23 | 23 | 1 | 23 |  |  | 8.11 | 0.05 |
| Y32out.2P-OFF | 168 | 101 | 44 | 18 | 144 | 6 | 15.20 | 0.57 |
| Y32in.3P-ON | 269 | 175 | 38 | 134 | 133 | 2 | 12.84 | 0.44 |
| SW1 Outlier | 145 | 123 | 54 | 77 | 66 | 2 |  |  |
| SW1 Disordered | 116 | 52 | 24 | 23 | 93 |  |  |  |
| Y71out.0P-GEF | 52 | 52 | 4 | 46 | 6 |  | 9.94 | 0.28 |
| Y71out.2P-SP2-A | 27 | 21 | 7 |  | 27 |  | 21.57 | 0.84 |
| Y71in.2P-SP12 | 20 | 15 | 2 |  | 20 |  | 5.73 | 0.25 |
| Y71out.2P-SP2-B | 19 | 9 | 6 |  | 17 | 2 | 15.2 | 0.42 |
| Y71out.2P-BINDER | 13 | 5 | 4 |  | 13 |  | 9.94 | 0.42 |
| Y71in.3P-R | 97 | 83 | 22 | 66 | 29 | 2 | 14.07 | 0.56 |
| Y71in.3P-SP12-A | 31 | 16 | 6 | 1 | 30 |  | 23.07 | 0.25 |
| Y71out.3P-T | 20 | 20 | 1 | 20 |  |  | 15.2 | 0.26 |
| Y71in.3P-SP12-B | 12 | 12 | 1 |  | 12 |  | 19.95 | 0.52 |
| SW2 Outlier | 250 | 176 | 79 | 104 | 145 | 1 |  |  |
| SW2 Disordered | 180 | 99 | 41 | 38 | 137 | 5 |  |  |

N=chains; PDB=entries; CF=crystal forms; NN=nearest neighbor; CA=alpha carbon; RMSD=root-mean-square deviations

**Supplementary Table S3.** Distribution of SW1 and SW2 conformations by α4α5 homodimerization status.

| Conformation Label | Homodimer | None | All |
| --- | --- | --- | --- |
| Y32out.0P-GEF |  | 23 | 23 |
| Y32out.2P-OFF | 20 | 148 | 168 |
| Y32in.3P-ON | 75 | 194 | 269 |
| SW1 Outlier | 20 | 125 | 145 |
| SW1 Disordered | 29 | 87 | 116 |
| Y71out.0P-GEF |  | 52 | 52 |
| Y71out.2P-SP2-A |  | 27 | 27 |
| Y71in.2P-SP12 |  | 20 | 20 |
| Y71out.2P-SP2-B |  | 19 | 19 |
| Y71out.2P-BINDER | 3 | 10 | 13 |
| Y71in.3P-R | 22 | 75 | 97 |
| Y71in.3P-SP12-A |  | 31 | 31 |
| Y71out.3P-T | 20 |  | 20 |
| Y71in.3P-SP12-B |  | 12 | 12 |
| SW2 Outlier | 50 | 200 | 250 |
| SW2 Disordered | 49 | 131 | 180 |
| All | 144 | 577 | 721 |

**Supplementary Table S4.** Distribution of SW1 and SW2 conformations by inhibitor chemistry.

| Conformation Label | SP12 |  |  |  | SP2 |  |  | Other | Multiple | None | All |
| --- | --- | --- | --- | --- | --- | --- | --- | --- | --- | --- | --- |
|  | IND | BEN | BIP | UNCL | ACR | SUL | UNCL |  |  |  |  |
| Y32out.0P-GEF |  |  |  |  |  |  |  |  |  | 23 | 23 |
| Y32out.2P-OFF | 8 |  |  | 8 | 47 | 4 | 13 |  | 2 | 86 | 168 |
| Y32in.3P-ON | 15 | 21 | 12 | 5 |  |  | 1 |  | 3 | 212 | 269 |
| SW1 Outlier | 4 |  |  | 5 | 3 | 1 | 9 | 4 |  | 119 | 145 |
| SW1 Disordered |  | 3 | 4 |  | 3 | 9 | 3 | 3 |  | 91 | 116 |
| Y71out.0P-GEF |  |  |  | 1 |  |  |  | 3 |  | 48 | 52 |
| Y71out.2P-SP2-A |  |  |  |  | 17 |  | 9 |  | 1 |  | 27 |
| Y71in.2P-SP12 | 4 |  |  | 4 |  |  |  |  |  | 12 | 20 |
| Y71out.2P-SP2-B |  |  |  |  | 6 |  |  |  |  | 13 | 19 |
| Y71out.2P-BINDER |  |  |  |  |  |  |  |  |  | 13 | 13 |
| Y71in.3P-R | 7 |  |  | 4 |  |  |  |  |  | 86 | 97 |
| Y71in.3P-SP12-A | 2 | 11 | 1 |  |  |  |  |  |  | 17 | 31 |
| Y71out.3P-T |  |  |  |  |  |  |  |  |  | 20 | 20 |
| Y71in.3P-SP12-B |  | 3 | 4 |  |  |  |  |  |  | 5 | 12 |
| SW2 Outlier | 6 |  | 1 | 8 | 25 | 12 | 13 | 1 |  | 184 | 250 |
| SW2 SDisordered | 8 | 10 | 10 | 1 | 5 | 2 | 4 | 3 | 4 | 133 | 180 |
| All | 27 | 24 | 16 | 18 | 53 | 14 | 26 | 7 | 5 | 531 | 721 |

## SUPPLEMENTARY DATASETS

**Supplementary Dataset S1.** List of RAS structures in the PDB annotated by SW1 and SW2 conformations and molecular contents.

**Supplementary Dataset S2.** Y32(OH):G12(CA) and Y71(OH):V9(CA) distances across RAS structures in the PDB.

**Supplementary Dataset S3.** Druggability scores and pocket volumes of inhibitor-bound and inhibitor-unbound sites across RAS structures in the PDB.

**Supplementary Dataset S4.** Y32(OH):3P(O1G) distances across 3P RAS structures in the PDB.

